# *FHY3*/*FAR1* transposable elements generate adaptive genetic variation in the *Bassia scoparia* genome

**DOI:** 10.1101/2023.05.26.542497

**Authors:** Nathan Hall, Jacob Montgomery, Jinyi Chen, Christopher Saski, Maor Matzrafi, Phil Westra, Todd Gaines, Eric Patterson

## Abstract

A near complete genome assembly consisting of 14 scaffolds, a total length of 969.6 Mb, and N50 scaffold length of 99.88 Mb, was generated to better understand how transposable element activity has led to adaptive evolution in *Bassia scoparia* (kochia), an agronomically important weed. The 9 largest scaffolds correspond to the 9 chromosomes of the close relative, *Beta vulgaris*. From this assembly, 54,387 protein-coding gene loci were annotated. We determined that genes containing Far-Red Elongated Hypocotyl 3 (FHY3) or Far-Red Impaired Response 1 (FAR1) functional domains have undergone a large, kochia-specific gene family expansion. We discovered that putative Mutator Don-Robertson (MuDR) transposable elements with detectable FHY3/FAR1 domains were tightly associated with segmental duplications of *5-enolpyruvylshikimate-3-phosphate synthase* subsequently conferring resistance to the herbicide glyphosate. Further, we characterized a new MuDR subtype, named here as “Muntjac”, which contributes to the evolution of herbicide resistance in kochia through the process of transduplication. Collectively, our study provides insights into the role of a *FHY3/FAR1* gene as an active transposable element and contributes new perspectives on the interaction between transposons and herbicide resistance evolution.

**Significance Statement:** In this work we discover that genes known as *FHY3/FAR1*, supposed “domesticated transposons”, are still actively transposing in the weed *Bassia scoparia* rearranging the genome in new ways. One rearrangement they are involved in is the tandem duplication of the gene *5-enolpyruvylshikimate-3-phosphate synthase* which is the gene that produces the target protein of glyphosate, one of the most important herbicides in the world. This work gives a specific example of the role that transposable elements play in making novel genetics and therefore the role that new genome rearrangements play in adaptative evolution, especially in response to external stress such as herbicides.

## 1. Introduction

Transposable elements (TEs) are engines of genetic change that can fundamentally alter an organism’s genome and genetic diversity, even within relatively short evolutionary timescales (Lisch 2013). TEs are active and mobile in the genome and propagate through replication cascades that are independent from the duplication of the rest of the genome. This process is primarily thought of as detrimental for overall organism fitness, as it can disrupt proper gene function and expression; however, TE activity can also occasionally generate adaptive alleles that contribute to novel, adaptive traits at the population or species level. The two major classes of TEs are defined by their propagation mechanism and classified as either Class I or II (Finnegan 1989). Class I TEs replicate via an RNA intermediate, utilizing a so-called “copy and paste” mechanism, while Class II TEs are without RNA intermediates and instead mobilize DNA through excision and re-insertion, via a “cut and paste” mechanism. When TEs insert into a new location in the genome, they may impart local structural changes to the DNA, such as methylation, that can have implications for gene expression for nearby genes (Burgess et al. 2020, Lisch 2009). At a chromosomal scale, TE insertions can facilitate large-scale genomic rearrangements such as deletions, inversions, and segmental duplications (Bennetzen 2000, Devos et al. 2002). At the gene scale, TEs can cause loss of gene function or gene shuffling through the process of transduplication (Candela and Hake 2008, Hsing et al. 2007, Juretic et al. 2005). Finally, and most rarely, TEs may contribute novel genes to the genome through a process of TE domestication, as in the case of the genes *far-red elongated hypocotyl 3* (*FHY3*) and *far-red-impaired response 1* (*FAR1*). These domesticated TEs were originally derived from a Class II TE known as a Mutator-like (Mule) transposon in the Mutator Don Robertson (MuDR) TE clade, but they have been hypothesized to have lost their ability to transpose and neofunctionalized as transcription factors that coordinate response to light signals (Dupeyron et al. 2019).

All functioning, autonomous MuDR elements have a shared set of characteristics: 1) a DDE motif in the transposase domain, 2) terminal inverted repeats (TIRs), and 3) target site duplications (TSDs) that are created when the MuDR is inserted in a novel location (Dupeyron, et al. 2019). When Mule elements transpose, including MuDRs, their excision and reinsertion is known to pick up a ‘payload’ of genomic material composed of DNA that originally flanked the element (Lisch 2015). The transposase domain of MuDRs has been well characterized and is mechanistic in the complete excision of the forward and reverse DNA strands from the genome, frequently leaving an excision footprint (Wells and Feschotte 2020). MuDR TIRs aid in the formation of their circularized double stranded DNA intermediate during transposition. TSDs form at the site of MuDR insertion and are typically short (8-11 bp) directed repeats adjacent to the TIRs (Wells and Feschotte 2020). *FHY3* and *FAR1* are notable in that as part of their domestication process, they have lost their TIRs, resulting in the prediction that *FHY3* and *FAR1* cannot form the DNA intermediate needed for transposition of Class II TEs (Lisch et al. 2001). An undomesticated TE, named *Jittery*, is sister to the *FHY3* and *FAR1* genes and retains its TIRs and mobility (Xu et al. 2004), suggesting that *FHY3* and *FAR1* could also be transpositionally active, with the presence of TIRs.

*Bassia scoparia* (L.) *Schrad*. (kochia) is a summer annual weed native to eastern and central Europe as well as western Asia (Friesen et al. 2009). It has become an economically important weed in crop and ruderal areas across the Canadian Prairies and the Great Plains of the USA (Kumar et al. 2019). Kochia is diploid with a chromosome number and genome size of 1n=9 and 0.7-1 Gb, respectively (Friesen, et al. 2009, Patterson et al. 2019). Glyphosate resistance in kochia is widespread and due to a tandemly arrayed copy number variation (CNV) of glyphosate’s target’s gene, *5-enolpyruvylshikimate-3-phosphate synthase* (*EPSPS*) (Jugulam et al. 2014, Wiersma et al. 2015). In previous work, an unidentified mobile genetic element (MGE) that contained a gene annotated as having FHY3 and FAR1 protein domains was identified flanking the *EPSPS* CNV, suggesting that the origin of the *EPSPS* CNV was closely related with *FHY3* and *FAR1* mobility (Patterson, et al. 2019). In this paper, we investigated an expansion of the *FHY3/FAR1-like* gene family in the kochia lineage and provide evidence that some of these *FHY3/FAR1-like* genes are not domesticated but are still actively mobile in the kochia genome. Furthermore, we specifically describe the exact FHY3/FAR1 MGE involved in glyphosate resistance and characterized it in the broader context of the MuDR transposon clade. We renamed this specific *FHY3/FAR1* gene associated with duplication of *EPSPS* “Muntjac”, after a highly mobile species of deer, to differentiate it from the domesticated *FHY3/FAR1* genes. We show evidence that *Muntjac* is mobile in the kochia genome and evidence of *FHY3/FAR1*gene family contraction and/or expansion in recent timescales. Furthermore, we hypothesize that the activity of *Muntjac* could be a signature of kochia’s genome plasticity as it continues to respond to agriculture-mediated selection.

## 2. Results

### 2.1. The kochia genome

A collapsed haploid reference genome was assembled using a combination of high coverage RSII PacBio single-molecule real-time sequencing (130 Gbp of raw data), deep-coverage Illumina shotgun sequencing (144 Gbp of raw data), a Hi-C linkage dataset (87 Gbp of raw data), and BioNano optical maps produced from an herbicide-sensitive plant. The assembly produced 14 scaffolds with a contig N50 length of 0.91 Mb and a scaffold N50 length of 99.88 Mb. Total sequenced length of the assembly was 837.60 Mb with 132.0 Mb in gaps (length of gaps estimated by optical maps). The total length of the assembly closely matched the haploid genome size estimated from flow cytometry 920-1070 Mbp (Supplemental Table S1). The nine largest scaffolds, corresponding to the nine chromosomes of kochia (Supplemental Figure S1), were ordered and numbered as pseudochromosomes based on synteny to the *Beta vulgaris* genome (Supplemental Figure S2). Due to *Beta vulgaris* and kochia being significantly phylogenetically divergent, several inversions, duplications and deletions are apparent in the whole genome alignment (Supplemental Figure S2). The five remaining, much smaller scaffolds from kochia remain unincorporated in the chromosome assembly due to a lack of information placing them in the larger pseudomolecules, although the two largest unplaced scaffolds appear to be non-collapsed (i.e. phased) regions of chromosome 3 (Supplemental Figure S2). We identified 95.0% of the 1,614 benchmarking universal single-copy orthologs (BUSCOs) in the embryophyta_odb10 dataset as either complete or fragmented in the genome assembly, which indicated a high degree of completeness for the gene space in the assembly (Supplemental Table S2). In total, 54,387 gene models were predicted; 45,372 received functional annotation; 8,333 unique Interpro annotations were assigned to 30,046 genes; and 2,234 unique Gene Ontology terms were assigned to 19,357 genes.

### 2.2. Gene family expansions and the FHY3/ FAR1-like gene family in kochia

Orthologous gene clustering using CAFÉ and all annotated genes from kochia and eight closely related species from the Caryophyllales clade (Chenopodiaceae and Amaranthaceae) revealed 157 gene families that are significantly expanded and 24 families significantly contracted in kochia relative to all other chenopods (Figure 1). Of the expanded gene families, five gene families contain genes annotated with FHY3/FAR1 family and/or FAR1 DNA binding domains (IPR031052 and IPR004330 respectively), accounting for 72 total genes. Furthermore, using the results from Orthofinder and MCSCANX, we see an especially high enrichment of *FHY3* and *FAR1* related genes (IPR004330, IPR031052, and/or IPR037766) physically linked with segmental duplications - a total 80 of the 282 detected segmental duplications contained at least one *FAR1* related gene that had undergone duplication as well. These duplications seem to be concentrated in gene rich areas of the chromosomes rather than the centromere and/or telomere (Figure 2).

**Figure 1.**
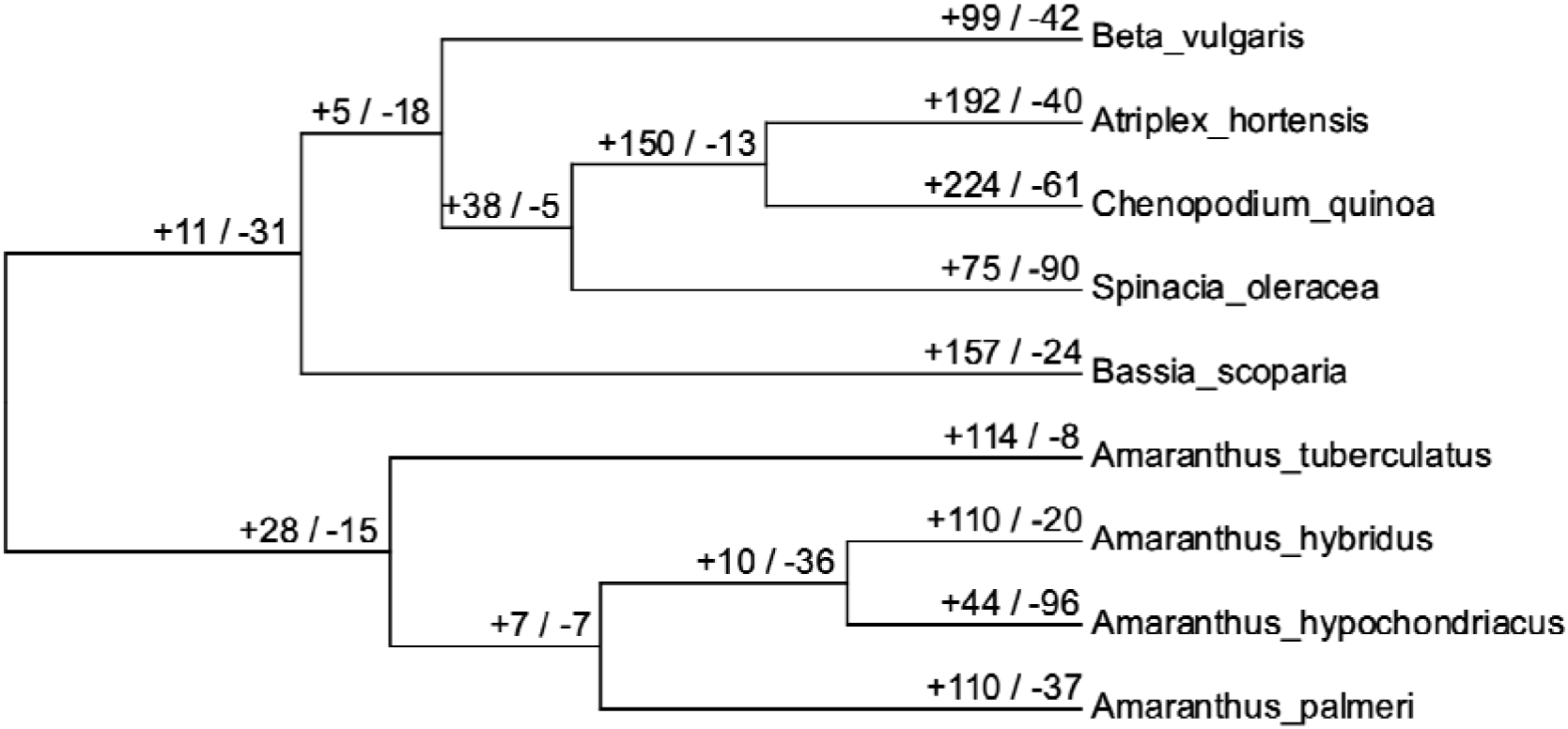
A cladogram of *Bassia scoparia* and several other *Amaranthaceae* species that have sequenced genomes. Numbers of significant gene family expansions (+) and contractions (-) (alpha=0.05) are listed along each branch as calculated by Computational Analysis of gene Family Evolution (CAFE).

**Figure 2.**
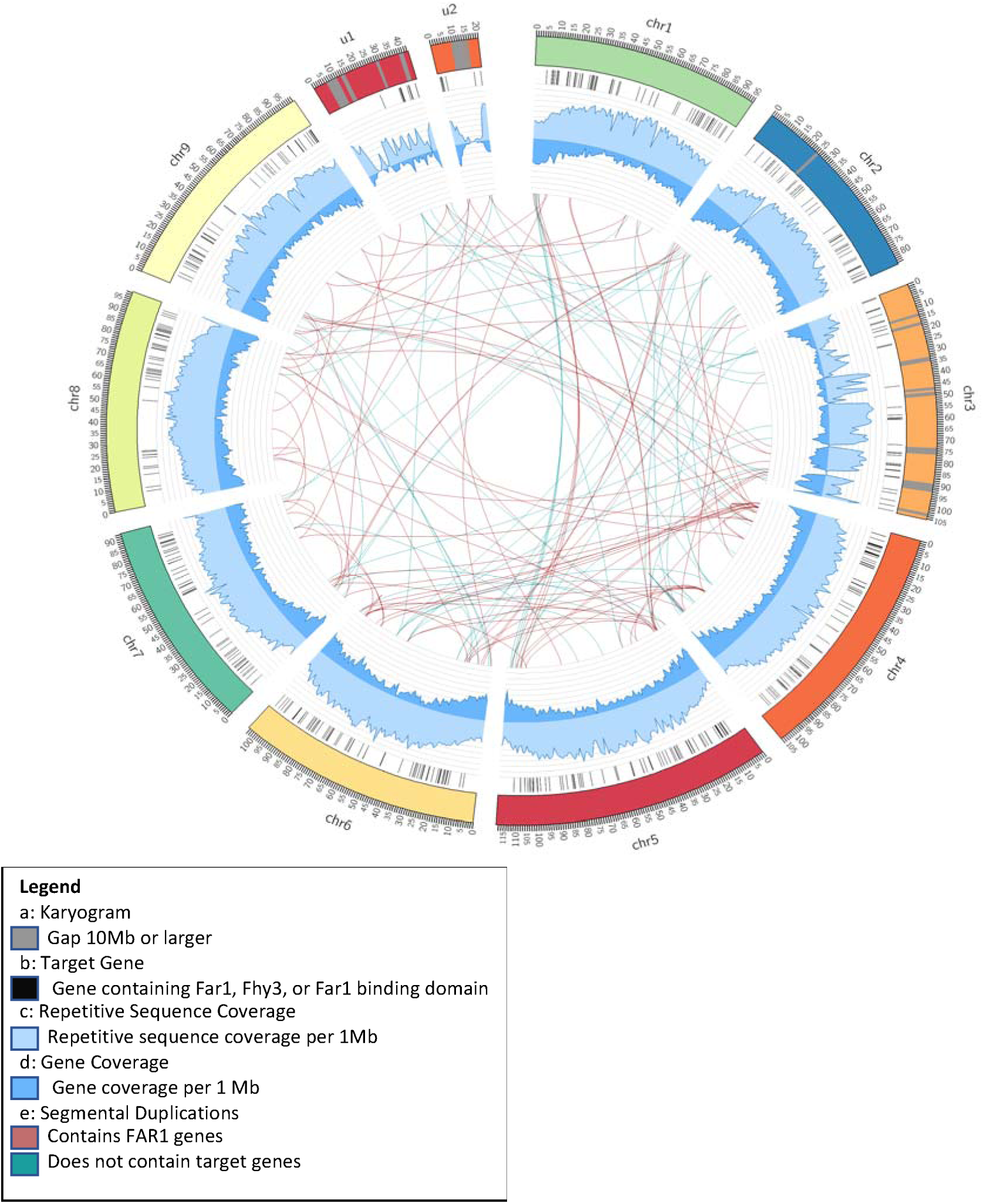
Circos plot of *Bassia scoparia* (kochia) genome assembly; a) chromosomes (chr1-chr9) and the two largest unplaced scaffolds (u1,u2) with gaps larger than 10 Mb indicated in gray; b) the placement of genes containing at least one of the following Interpro designations: *FHY1* (IPR037766), *FAR1* DNA binding domain (IPR004330), and *FHY3*/ *FAR1* family (IPR031052); c) repetitive sequence coverage per 1 Mb; d) gene coverage per 1 Mb; e) segmental duplications detected using MCScanX. Red lines indicate segmental duplications that contain a *FAR1* gene. Green lines represent segmental duplications that do not contain a *FAR1* gene.

### 2.3. Copy number variation between glyphosate-resistant and -sensitive plants

While several different arrangements of the *EPSPS* CNV have been suggested (Ravet et al. 2021), the *EPSPS* CNV event being described here is confirmed to: 1) be in a tandem arrangement (i.e., not dispersed) with two distinct repeat units of different sizes, 2) include seven genes (one of which was *EPSPS*), and 3) be associated with the insertion of one or more ∼15 kb MGEs containing the complete *Muntjac* gene. This specific arrangement was previously confirmed in the glyphosate-resistant line, M32, by bacterial artificial chromosome sequencing (Patterson, et al. 2019). These MGEs are flanked by TIRs suggesting that the MGE includes *Muntjac* (∼3 kb) plus additional genetic payload between the TIRs.

Shotgun Illumina resequencing from a glyphosate-resistant plant identified dozens of small CNV events between the resistant and sensitive samples, but none so large or nearly as high copy number as the *EPSPS* CNV event. The MGE is present as a single copy in the susceptible genome on chr9, and this MGE sequence from the susceptible genome is nearly identical to the MGE located next to *EPSPS* on chr1 in the resistant line (>99.9% sequence similarity). Breaks in read mapping coverage indicate the presence of a structural variant between the MGE at chr9 in the susceptible plant and the MGE on chr1 in the resistant plant (Figure 3).

**Figure 3.**
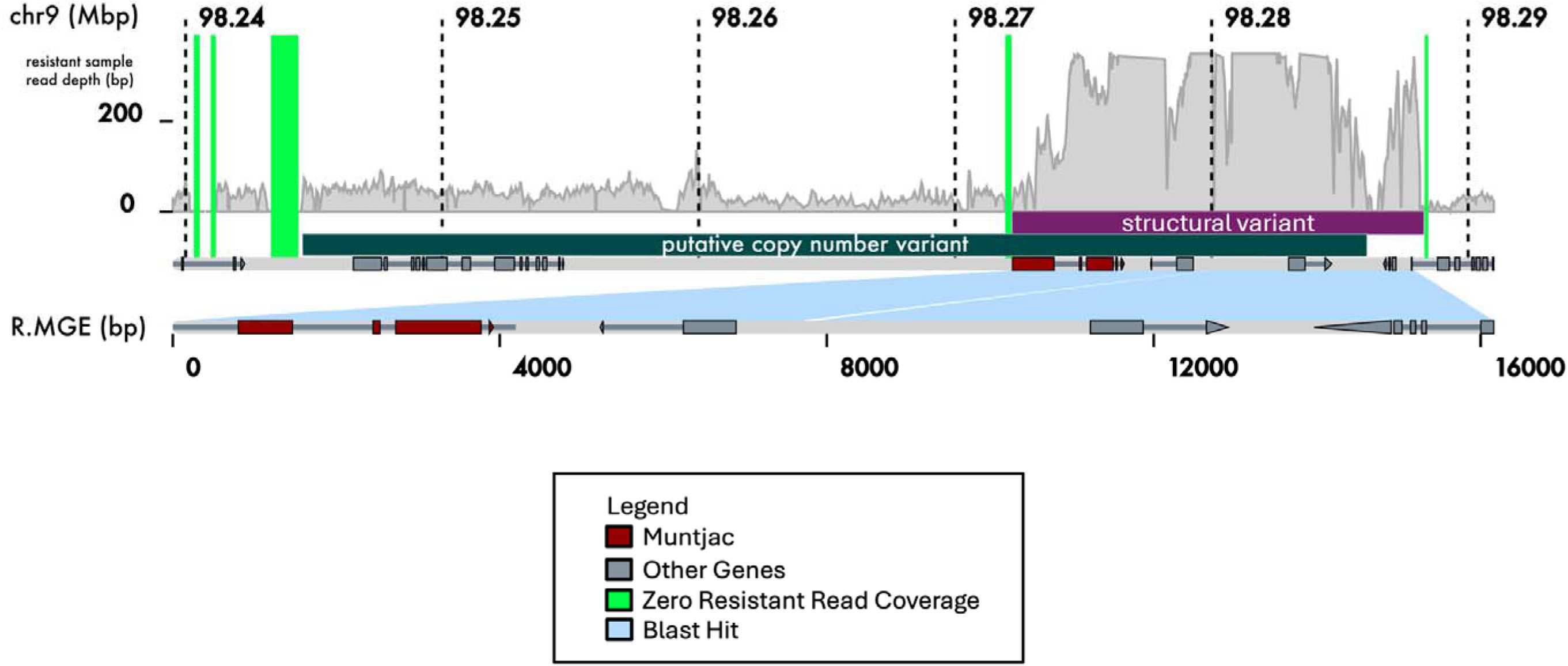
Coverage of genomic DNA Illumina reads from a glyphosate-resistant *Bassia scoparia* (kochia) plant mapped to the genome assembly of a sensitive nt. Putative copy number variants predicted using CNVnator. Structural variants predicted using complete breaks in read mapping, and micro-assembly.

### 2.4. Muntjac activity and transposition

The results of CNVnator and read mapping indicate that *Muntjac* is within the MGE that is present in the CNV related to glyphosate resistance in kochia. Interruptions and variations in read coverage are due to the MGE containing *Muntjac* that occurs at different loci within the glyphosate resistant population, consistent with transposase activity (Figure 3). Micro-assemblies of the susceptible chr9 and resistant chr1 insertion sites of the MGE provide further support for *Muntjac* as an autonomous Mutator transposase (Figure 4). First, a TSD was detected at both the insertion next to *EPSPS* on chr1 (∼6,110,000 bp) in the resistant line as well as at its location on chr9 (∼98,280,000 bp) in the assembled susceptible line. Second, a TIR of the appropriate length (88 bp) was identified adjacent to each TSD and flanking both sides of *Muntjac*. A survey of protein domains within *Muntjac* and other MuDR type demonstrates that *Muntjac* retains key conserved domains for transposase activity including a Mutator domain (MULE) and a SWIM zinc-finger (Figure 5). The detection of a putative YCF1 (Yeast Cadmium Factor-1) domain within the Muntjac protein is unique and does not appear to be in any other *FHY3* or *FAR1* genes in the susceptible genome. It should be noted that we also identified another putative *Muntjac* copy at a novel position on chr1 (∼22,725,000 bp) in the genome assembly; however, it was missing TIRs and had fused with another unrelated gene, suggesting that this copy is not transposable (Figure 4).

**Figure 4:**
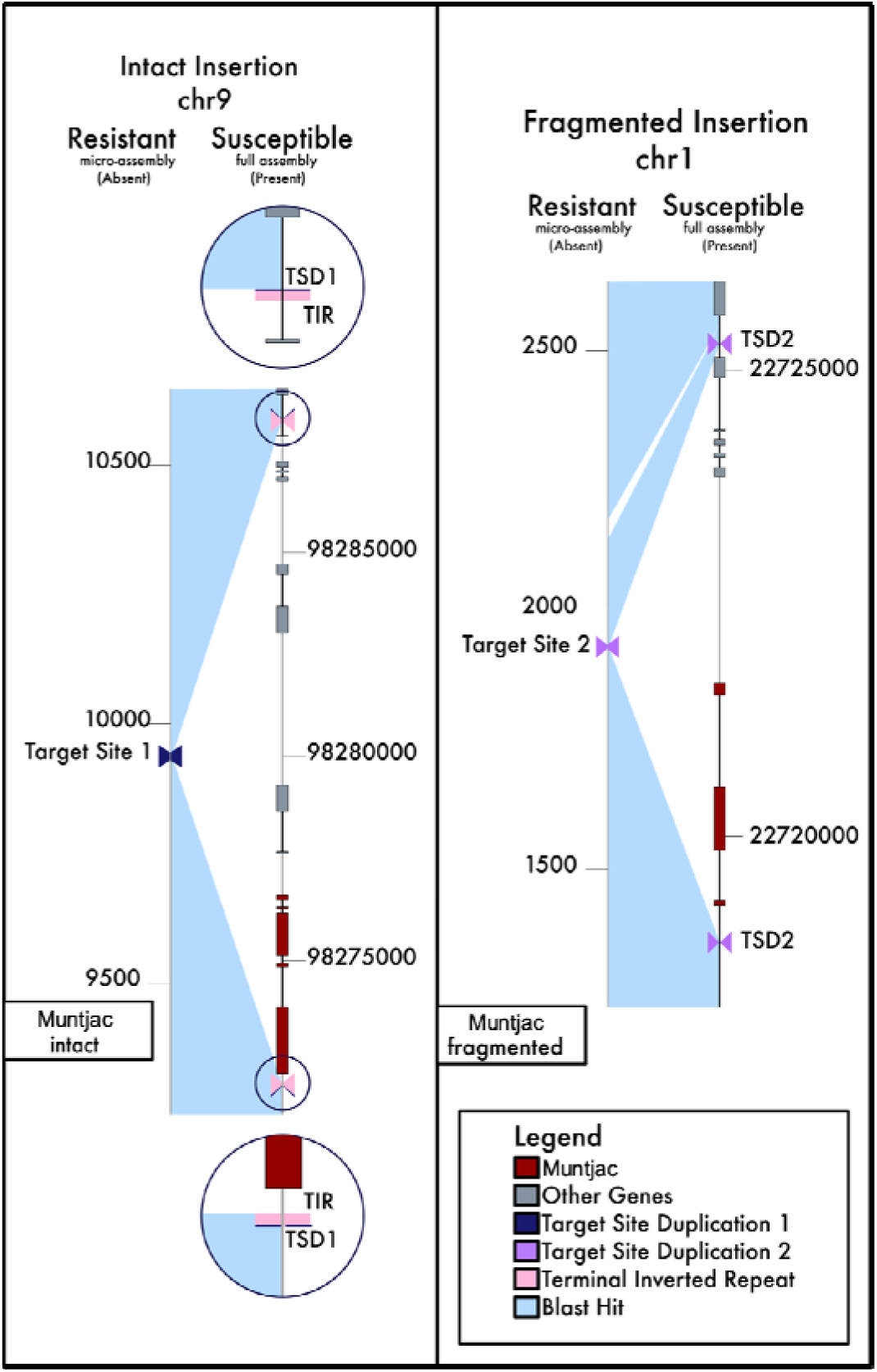
Micro-assemblies from the glyphosate-resistant *Bassia scoparia* (kochia) line (M32) were produced for two *Muntjac* insertion sites that were detected in the susceptible line (7710). a) A completely intact Muntjac insertion on Chromosome 9, with detected target site duplication (TSD), and detectable terminal inverted repeats (TIR) that is present in the susceptible assembly, but not the resistant. b) A fragmented insertion on Chromosome 1 that is also present in the susceptible but not resistant individual. The *Muntjac* gene has been pseudogenized or incorporated into another a gene in the susceptible individual. The TSD provides evidence of insertion, but the fragmentation of *Muntjac* and the absence of TIRs suggest that this insertion on Chromosome 1 is no longer active.

**Figure 5:**
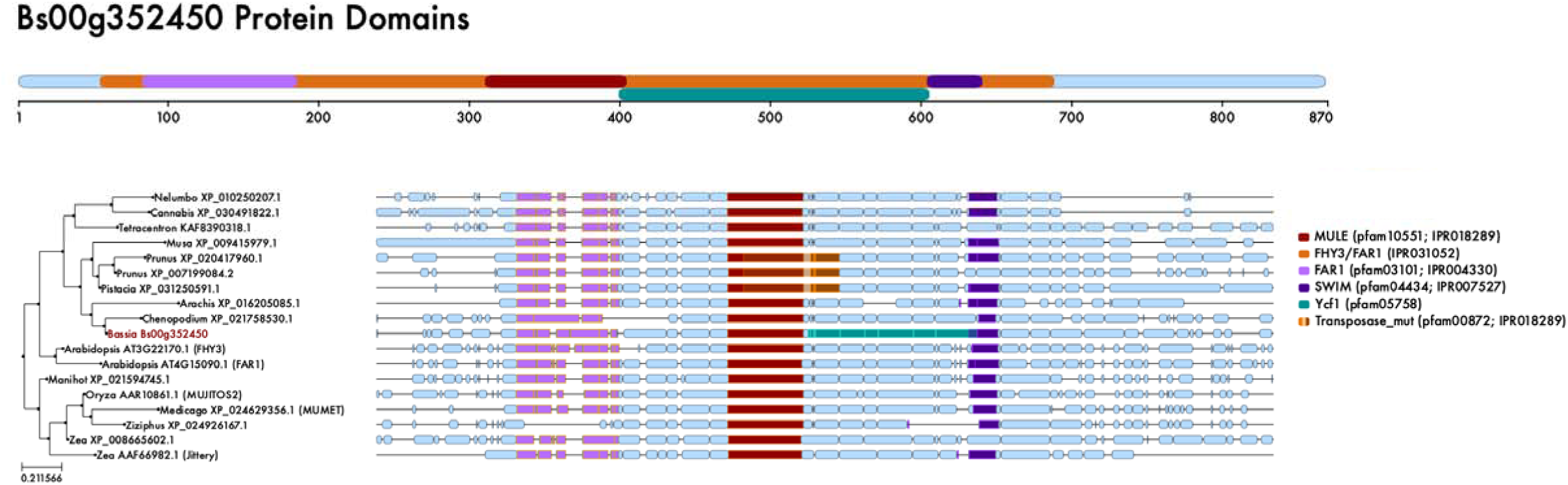
A gene tree produced from an alignment of representative MuDR sequences from *Bassia scoparia* (top track) and several other plant clades showing ctional domains within the MuDR clade. The *FHY3*/*FAR1* domain was detected in all sequences but is not shown for clarity.

### 2.5. Molecular verification of Muntjac activity and transposition

Active MuDR elements are mobile and therefore can be found at different loci in different individuals. To determine if *Muntjac* exhibited mobility within the kochia genome, we designed five pairs of PCR primers (Supplemental Table S3) that detect the absence/presence of *Muntjac* at chr9 (∼98,280,000 bp) in a variety of kochia plants from diverse locales and with varying levels of glyphosate sensitivity (Figure 6, Table 1). These primers were tested and validated using DNA from known glyphosate-sensitive and -resistant populations (named 7710 and M32, respectively) as we have genomic information supporting the expected presence/absence of amplification in these plants. As predicted, the F1/R1 primer pair did not amplify for samples from 7710 since the MGE fragment (∼16,000 bp) was too long to be amplified under the PCR setup. In contrast, DNA from M32 plants produced the predicted ∼900 bp target band, indicating the MGE was not in its chr9 position in this resistant line. Primer pairs F1/R4 and F3/R3 amplified the junction between MGE and the insertion point, which further demonstrated the presence of the full MGE on chr9. As expected, 7710 plants showed target bands with both primers while M32 plants did not. Primer pairs F1/R2 and F2/R3 were used to verify that the assembled DNA sequence flanking the chr9 insertion site was present and correct (Figure 6, Table 1).

**Figure 6:**
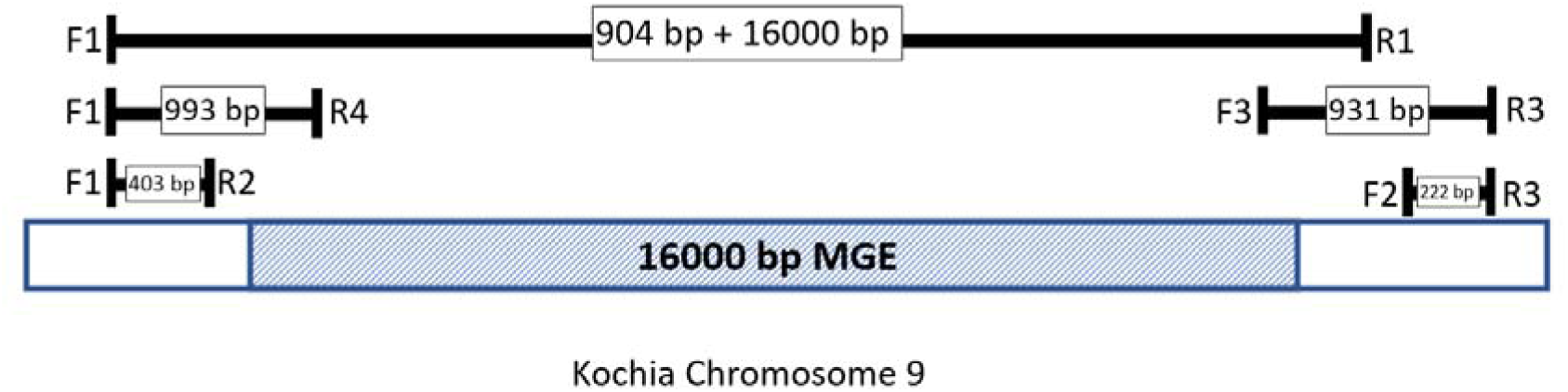
Diagram of the primer positions used to verify the presence/absence of MGE sequence on Chromosome 9 in glyphosate-resistant and -susceptible *Bassia scoparia* (kochia) individuals.

**Table 1.** Results of MGE validation primer pairs.

| Assay | Primer pair | Populations |  |  |  |  |  |  |  |  |  |
| --- | --- | --- | --- | --- | --- | --- | --- | --- | --- | --- | --- |
|  |  | Oregon |  | CO-S |  | Israel |  | M32 |  | 7710 (S mixture) |  |
|  |  | Oregon-1 | Oregon-2 | CO-S-1 | CO-S-2 | Israel-1 | Israel-2 | M32-1 | M32-2 | S-1 | S-2 |
| 1 | F1/R1 | × | ✓ | × | ✓ | ✓ | ✓ | ✓ | ✓ | × | × |
| 2 | F1/R2 | ✓ | × | ✓ | ✓ | ✓ | ✓ | ✓ | ✓ | ✓ | ✓ |
| 3 | F2/R3 | ✓ | × | ✓ | ✓ | ✓ | ✓ | ✓ | ✓ | ✓ | ✓ |
| 4 | F1/R4 | × | × | × | × | × | × | × | × | ✓ | ✓ |
| 5 | F3/R3 | × | × | × | × | × | × | × | × | ✓ | ✓ |

✓: band of/close to target length amplified
×: band of/close to target length not amplified
\* Multiple bands amplified

We also screened novel kochia populations with and without increased *EPSPS* copy number but no other genomic information, to determine if the MGE containing *Muntjac* is present on chr 9, as it is in the glyphosate-sensitive population, 7710. We selected a glyphosate-resistant line from Oregon and two susceptible populations (CO-S and a population collected from Israel in the native range of kochia). In these populations, plants Oregon-1, CO-S-1, Israel-1, and Israel-2 showed target length bands with primer F1/R1, suggesting the MGE is not present at the chr9 locus in any of these resistant and susceptible populations. As anticipated from this result, the primer sets used to test the boundaries of the MGE (F1/R4 and F3/R3) also did not amplify (Table 1). Amplification with primer pairs F1/R2 and F2/R3 confirmed that the sequence around the chr9 insertion was conserved across accessions (Table 1). Therefore, in these plants, the MGE is not present at the chr9 locus where it is found in the 7710 population. For plants Oregon-1 and CO-S-1, there were target bands produced by F1/R2 and F2/R3, but three primer pairs (F1/R1, F1/R4, and F3/R3) used to detect the MGE presence/absence did not amplify to confirm its presence or absence. In Oregon-2, there was no MGE insertion at chr9, as F1/R1 produced a band, but F1/R4 and F3/R3 did not. However, in this individual, multiple bands were identified with primers F1/R2 and F2/R3. Results in this plant suggest the presence of sequence variations in this region among different kochia individuals. Overall, the primer pairs used for validation here can be applied to most kochia populations and have demonstrated that MGE position varies with kochia plants/populations. We conclude that the MGE at the chr9 position is not shared among all susceptible populations and seems to be its location only in the ‘7710’ reference genome.

## 3. Discussion

### 3.1. The Kochia Genome

Chromosome-scale reference genomes are critical for investigating weed genomics and evolution questions (Montgomery et al. 2024a). The genome of *Bassia scoparia* comprises approximately 1 Gb of genetic material, which is characterized by high repetitive element content (Supplemental Table S4), similar to other chenopod species. Through an integrative approach that employed short- and long-read sequencing techniques, chromatin contact mapping, and optical mapping we achieved an assembly that is nearly the size predicted through flow cytometry (∼97%). The assessment of gene space completeness using the embryophyte odb10 dataset via BUSCO estimates was found to be 95.0%. Although the development of newer sequencing and assembly technologies may allow for a more contiguous kochia genome, the assembly’s comprehensive coverage of gene space and relevant *EPSPS* and *Muntjac* loci allowed us to accomplish our objective of understanding *Muntjac* mobility across kochia populations. In fact, this reference genome is already being successfully used to map other transposon-related herbicide resistance traits (Montgomery et al. 2024b).

*Beta vulgaris* is the closest related species to kochia that has a chromosome-level genome assembly. Like most chenopods, both species have nine chromosomes. Alignment of the two genomes demonstrates a significant degree of synteny and collinearity between the species (Supplemental Figure S2); however, owing to their evolutionary divergence, numerous inversions, gaps, and transpositions have occurred. It may be that misassembles of either species also account for some of these differences. Furthermore, available data indicates that the *Beta vulgaris* genome is estimated to encompass approximately 758 Mb, approximately 75% of kochia’s genome size. The basis of this discrepancy is not yet apparent; however, preliminary analysis indicates the presence of a high number of transposable elements (TEs) in kochia (Supplemental Table S4). Notably, there is currently no evidence of any polyploidy events or significant genomic duplication in either species since their divergence.

### 3.2. FHY3/ FAR1-like gene family in kochia

When the annotated genes of kochia are compared to other sequenced chenopods and amaranths, there are several gene families that are significantly expanded in kochia according to our CAFE analysis (Figure 1). Many of these enriched families contain FHY3/FAR1 domains (Figure 2) and/or have functions related to TEs (e.g. IPR005162: Retrotransposon gag domain, IPR001878: Zinc finger, CCHC-type superfamily, and IPR00752: Zinc finger, SWIM-type). Past research indicated that one of these *FHY3/FAR1* genes was in a MGE that caused a CNV of the gene *EPSPS* (Patterson, et al. 2019). This CNV, in turn, endows resistance to the herbicide glyphosate and is the major mechanism by which kochia has adapted resistance to this herbicide (Gaines et al. 2016, Wiersma, et al. 2015). *FHY3/FAR1* genes are rare cases of ‘domesticated’ transposons. They have supposedly lost their ability to readily translocate and self-duplicate, but they have been co-opted as transcription factors (Hardigan et al. 2016, Lisch, et al. 2001). The expansion of *FHY3/FAR1* genes may indicate that they have either 1) retained their transposable element capabilities or 2) they have re-acquired transposable element capabilities in the kochia lineage.

The interaction between genomic rearrangements and adaptation underscores the profound impact that genomic plasticity can have on plant evolution, particularly under sustained abiotic stresses (Johnson et al. 2025, Sen et al. 2024). While not a novel concept, previous research has shown that TE activity and CNVs are essential pathways for generating novel genetic diversity. Nevertheless, the potential contribution of structural variation as a driver of adaptive genetic variation is often underestimated as supported by several studies (Fuentes et al. 2019, Horváth et al. 2017). TEs contribute to genetic diversity by generating a wide range of mutations, which can help mitigate genetic bottlenecks and facilitate rapid responses to novel conditions (Casacuberta and González 2013). For instance, in the invasive weed *Mikania micrantha*, a noxious invasive weed found in southern China, researchers identified 59 genetic and 86 epigenetic adaptive TE loci, with 51 genetic and 44 epigenetic loci correlating with 25 environmental variables, supporting rapid adaptation and invasion under different climatic conditions (Su et al. 2021). In *Capsella rubella*, frequent TE insertions at *FLOWERING LOCUS C* account for 12.5% of the natural variation in flowering time, a critical life history trait linked to fitness and adaptation in changing new environments (Niu et al. 2019).

Gene duplication has the unique advantage of changing gene expression without changing the protein structure or regulatory elements controlling gene expression. This process is dynamic and responsive to abiotic stress, allowing selection to quickly increase or decrease average copy number within a population. Our study utilized deep-coverage, short read genomic sequences of a glyphosate resistant individual to examine the *EPSPS* locus and the *FHY3/FAR1* MGE described previously (Patterson, et al. 2019). A genome-wide survey was conducted to explore loci, beyond the *EPSPS* locus, that demonstrated variation in copy number within the resistant genome. As anticipated, given the sequence divergence between the two Kochia individuals, the analysis revealed hundreds of small copy number variants (CNVs) at both lower and higher copy number than the *EPSPS* locus. Many of the CNVs at very high read depths contained genes annotated as TEs.

### 3.3. Muntjac activity and transposition

*Muntjac*’s location within the genome is variable across several kochia samples. We designed PCR markers that amplify if, and only if, the MGE is at its position on chr9, as in the susceptible genome. Additionally, we developed markers that amplify only if the MGE is absent from this locus (Figure 6). The former primers span the insertion boundaries and only amplify if the MGE is present on chr9. The latter primers span the upstream and downstream boundaries of the MGE and the surrounding genome. If the entire MGE is present, the amplicon is too large to amplify (∼16 Kb); but if it is absent, it is only 900 bp. We ran these molecular markers on the susceptible kochia line from which the genome was developed as well as from the glyphosate resistant line from which we first identified *Muntjac*. We confirmed that the *Muntjac*-containing MGE is indeed missing from its position on chr9 in the glyphosate resistant line, M32, while it is present in the glyphosate susceptible line, 7710. From these data we hypothesized that the *Muntjac*-containing MGE transposed from its position on chr9 to the *EPSPS* locus on chr1 using a “cut-and paste” mechanism. To test whether chr9 is indeed the “native” location for *Muntjac*, we used our markers on two unique susceptible lines (divergent to the genotype used to assemble the reference genome) and a glyphosate resistant line unrelated to the M32 glyphosate-resistant population. *Muntjac* was not at the position on chr9 or the position on chr1 (underlying resistance in concert with the *EPSPS* CNV), in any of these additional populations or individuals. This indicates that a conserved, original locus of *Muntjac* is still unknown, if it exists at all. Recently, *EPSPS* duplication without *Muntjac* as part of the CNV has been identified in some kochia populations, indicating multiple unique paths to *EPSPS* copy number variation (Ravet, et al. 2021). This suggests high levels of structural variation, and therefore genetic diversity, exist in the kochia species.

The *FHY3/ FAR1* gene in the MGE that caused the *EPSPS* CNV is a complete transposon, containing key protein domains for transposition. Close molecular examination of the MGE and *Muntjac* in a glyphosate-resistant line from Oregon shows evidence of differential transposition when compared to resistant and susceptible lines from Colorado (CO) and Israel. This line of evidence demonstrates the variability of the insertion site described from M32 on chr9. The detection of TSD, TIRs, and the apparent mobility of *Muntjac* provides strong lines of evidence supporting active transposition of *Muntjac* within kochia.

In summary, we assembled the genome of a glyphosate-susceptible kochia plant to chromosome-scale. By performing orthologous gene clustering of related chenopods and amaranths, we observed an expansion of genes containing FHY3 and/or FAR1 domains in kochia. Resequencing of a glyphosate resistant plant using high coverage Illumina data revealed the location of a MGE that is involved in the formation of the *EPSPS* CNV. Within this element, a gene annotated as having FHY3 and FAR1 protein domains was identified. Furthermore, this element had signature target-site duplications, terminal inverted repeats, and a functional transposase domain in the *FHY3/FAR1* gene. We have named this element “Muntjac” to distinguish it from other *Mules* and *FHY3/FAR1* related genes. We verified that *Muntjac* exists in at least two different locations across different individuals using molecular markers: chr9 (∼bp 98,280,000) in the glyphosate-susceptible individual we assembled and flanking the glyphosate target gene *EPSPS* on chr1 (∼6,110,000 bp) in a glyphosate-resistant individual. We used these markers to understand if the MGE on chr9 is present across diverse herbicide-sensitive lines (including plants from Israel, kochia’s native range) and did not find *Muntjac* at either location. We hypothesize that *Muntjac* is still active within kochia and that its recent activity may only be resolvable using a more in-depth pangenomic approach. Regardless, we believe *Muntjac*’s involvement with generating glyphosate resistance alleles is a strong demonstration of the critical role TEs play in the production of genetic variation, and therefore, the adaptive potential of weedy species.

## 4. Material and methods

### 4.1. Germplasm

Several collections of glyphosate-susceptible and -resistant kochia plants were used in this study. The herbicide-susceptible reference genome assembly was generated from the highly inbred line known as ‘7710’, originally collected from western Nebraska (Preston et al. 2009). This line has been used extensively as an herbicide-susceptible control for many studies. The 7710 line has a consistent phenotype and is well-studied compared to many kochia collections (Busi et al. 2018, Pettinga et al. 2018). The primary glyphosate-resistant line used for comparison was the inbred line known as ‘M32’, collected from north-eastern Colorado (Westra et al. 2019), which has been confirmed to have increase gene copy number of *EPSPS* and consistent inheritance of the glyphosate resistant phenotype (Gaines, et al. 2016, Patterson, et al. 2019). Furthermore, the tandemly-duplicated *EPSPS* locus in M32 has been extensively studied and assembled using BAC sequencing (Patterson, et al. 2019).

To compare to more diverse genotypes, we collected two more glyphosate-susceptible populations; one from Israel in the native range of kochia (referred to as Israel), and a second from northern Colorado (referred to as CO-S). Additionally, we included glyphosate resistant plants with high *EPSPS* copy number but no amplification of *Muntjac* (referred to as Oregon) (Ravet, et al. 2021). For all studies, seeds were sown in SureMix®, grown in growth chamber (25 °C in the day and 20 °C at night, 14/10 h light/dark period), and watered every other day.

### 4.2. Genome size estimation and chromosome counting

Young leaf tissue of kochia (accessions 7710 and M32) and corn was sent to the Flow Cytometry and Imaging Core laboratory at Virginia Mason Research Center for genome size estimation via flow cytometry using an internal standard (2.5 pg DNA/cell) co-chopped with each sample. Corn samples were included to ensure the method produced accurate results. To determine the number of chromosomes for kochia, DNA of root tip cells undergoing metaphase was stained and visualized as part of a fee-for-service agreement with the Stack lab at Colorado State University.

### 4.3. Kochia genome sequencing and assembly

DNA was extracted from leaf tissue of a single glyphosate-susceptible kochia plant from the line 7710 using a modified CTAB protocol (Doyle and Doyle 1987, Li et al. 2013, Patterson, et al. 2019). DNA was sent for PacBio Sequel sequencing at the University of Delaware DNA Sequencing & Genotyping Center. Expanding leaf tissue was sent to Phase genomics for HiC library prep and sequencing. An optical map was constructed at the McDonnell Genome Institute using the BioNano Genomics Saphyr system. Short whole-genome shotgun paired end reads (insert size ∼350 bp) were generated at the Roy J. Carver Biotechnology Center at the University of Illinois on an Illumina HiSeq.

Long read sequences were initially assembled using MeCat2 (Xiao et al. 2017). Short reads were then aligned to this assembly using HiSat2 v2.1.0 (Kim et al. 2019). The assembly and alignment were used in conjunction to polish the genome using Pilon (Walker et al. 2014). Phased haplotigs were merged using HaploMerger2 (Huang et al. 2017). Merged contigs were scaffolded using HiC data and the SALSA pipeline (Ghurye et al. 2017). After initial scaffolding, the assembly was further scaffolded using optical maps generated at the McDonnell Genome Institute with the BioNano Solve software (Staňková et al. 2016). This assembly was then gap filled using PBJelly (English et al. 2012). Newly filled gaps were polished once again using Pilon. Finally, super-scaffolds were manually scaffolded into pseudo-chromosomes using the HiC data and Juicer (Forcato et al. 2017).

Gene models in the assembly were predicted using the GenSAS pipeline (Humann et al. 2019). The genome assembly was uploaded to the GENSAS webserver along with RNA-seq data for annotation (Pettinga, et al. 2018). The RepeatMasker and RepeatModeler tools were used to identify the repeats and mask the genome for gene model calling (Flynn et al. 2020, Tempel 2012). Publicly available RNAseq data (NCBI SRA: SRX3276189-SRX3276192) were aligned to the masked genome using BLAST+, BLAT, HISAT2, TopHat, and PASA (Cock et al. 2015, Haas et al. 2008, Kent 2002, Kim, et al. 2019, Kim et al. 2013). The genome and these alignments were run through the Augustus, BRAKER, GeneMark-ES, Genscan, and SNAP tools for gene model prediction with default parameters (Hoff et al. 2016, Lomsadze et al. 2005, Stanke et al. 2004, Ter-Hovhannisyan et al. 2008). Next, we made a consensus set of gene models using EvidenceModeler. The following weights were assigned to each predictive tool: 10 for transcript alignments and BRAKER; 5 for Augustus and protein alignments; 1 for GeneMark-ES and SNAP. PASA refinement was conducted using BRAKER-HISAT2 as the gene set. For the functional annotation, predicted genes were aligned to the NCBI plant protein database using both BLASTp and Diamond. Targeting and subcellular localization was predicted using Pfam, SignalP and TargetP, with default parameters for plants. Additional functional annotation was performed with Multiloc2, Iprscan, and an MMseqs2 search of the Uniref50 database (Steinegger and Söding 2017, Suzek et al. 2007). The final annotation was tested for completeness by running BUSCO using the embryophyta_odb10 lineage database on the predicted proteins (Waterhouse et al. 2017). Regions of synteny between kochia and *Beta vulgaris* were determined using GENESPACE (Lovell et al. 2022, McGrath et al. 2023).

### 4.4. Gene Family Expansions and the FHY3/ FAR1-like gene family in kochia

To determine the history of *Muntjac* and if FHY3/FAR1 domains were prevalent in gene expansions, orthogroup clustering was performed using Orthofinder v2.4.0 (Emms and Kelly 2019). All annotated genes in the kochia genome were subjected to orthologous gene clustering with all annotated genes from eight closely related species within Amaranthaceae (Supplementary Table S5). The species tree was inferred by Orthofinder (Emms and Kelly 2019). A CAFE analysis was conducted to detect significant expansions within orthogroups (De Bie et al. 2006).

The FHY3/FAR1-like protein described by Patterson et al. (2019) was used to validate the annotation of *Muntjac* in the genome assembly. The gene sequence was identified by protein blast to the assembly, and reads from three different SRA accessions (ERR364385, SRR6165247, and SRR1198330) were mapped to the gene with HISAT2. Predicted splice junctions from these alignments were used to confirm correct gene structure. The manually produced transcript was searched for ORFs using NCBI ORF Finder, searched for conserved domains using NCBI Conserved Domain Search and using the InterPro online search portal.

To classify types of gene duplication and detect perturbations to genome structure, MCSCANX was run according to its documentation (Wang et al. 2012). To confirm structural differences between the glyphosate susceptible line, 7710, and glyphosate resistant line, M32, micro-assemblies were created for the genomic regions surrounding an intact *Muntjac* (Bs.00g352450) and a fragmented *Muntjac* insertion (Bs.00g145150). To do this, genomic DNA sequencing reads from a glyphosate-resistant plant were mapped against the kochia genome with HISAT2, and mapped reads from target genes and the surrounding sequence were extracted with SAMtools v1.10 (Li et al. 2009). Extracted reads were assembled with SPAdes v3.14.0 (Bankevich et al. 2012) Assembled regions from resistant reads were compared back to each locus of origin in the sensitive genome to determine if an excision had occurred and search for target site duplications and terminal inverted repeats (TIRs).

To determine the relationship of *Muntjac* to *FHY3*, *FAR1* and *Jittery*, a gene tree was produced using protein sequences that represented several TE families including Jittery, Hop, and MuDR. Briefly, full protein records were found on NCBI by searching for the *FHY3*/*FAR1* domain sequence in the NCBI nr database using BLAST. Amino acid sequences of several elements from various taxa were aligned and curated. The curated alignment was analyzed using IQTree v1.6.12 for Windows (Nguyen et al. 2015) with flags (-alrt 1000, -bb 1000). For the purposes of visualization, the set of proteins was searched for Pfam domains using NCBI Conserved Domain Search. Figures were prepared with Inkscape v0.92.4.

### 4.5. Identifying copy number variants

The resequencing data used to generate micro-assemblies (described in section 4.3) was also used to determine genome-wide copy number variation (CNV) in the previously characterized glyphosate resistant line M32 (Patterson, et al. 2019). Paired end data were aligned to the susceptible reference genome assembly (from population 7710) using HISAT2. Additionally, Illumina paired end data from the susceptible plant was also aligned with HISAT2 to the reference to control for shared CNV events between the resistant and susceptible plants compared to the reference assembly. The resulting alignments were scanned using CNVnator v0.4.1 and a 1,000 bp sliding window to detect read coverage changes outside of the normal read depth distribution (Abyzov et al. 2011). Duplicated and deleted regions were identified using a read depth cutoff of 2x and 0.5x coverage for duplications and deletions (respectively) and an adjusted p-value < 0.05. CNV regions shared between the susceptible and resistant lines were removed from consideration, as they do not differentiate populations and are therefore likely not subject to glyphosate selection.

### 4.6. Muntjac and its MGE activity and transposition

PCR primers were designed to identify 1) the upstream and downstream boundaries of the MGE containing *Muntjac*’s insertion on chromosome 9 (chr9), 2) the excision of the MGE from chr9 in a glyphosate resistant individual, 3) the insertion of the MGE at the *EPSPS* locus on chr1, and 4) the presence of the flanking genomic sequence at the chr9 position. For this assay, we used several individuals from the five kochia collections described in section 4.1: 7710 (S), CO-S (S), Israel (S), M32 (R), and Oregon (R). DNA was extracted from individual plants of all populations. There were two replicates per population. Primers were designed based on kochia chr9 assembled sequences and were blasted to the whole kochia genome to confirm their specificity. The five pairs of primers were used for validation (Supplemental Table S3). The PCR was conducted in a 25-μL volume, consisting of 1 μL of DNA, 1 μL of each primer (10 μM) and 12.5 μL of GoTaq Green Master mix (Promega®, USA). The PCR was run in a thermocycler (BioRad, USA) with the following profile: 95 °C for 5 min, 35 cycles of 95 °C for 30 s, 55 °C for 30 s, and 72 °C s for 1 min, followed by a final extension step of 5 min at 72 °C.

## Data availability

The assembled genomes, associated GFF annotation files, and all functional annotation information are publicly available through the International Weed Genomics Consortium online database, ‘Weedpedia’ (https://weedpedia.weedgenomics.org/). The genome assembly is also available through NCBI under accession SNQN00000000. Data used to generate the reference genome assembly is available through NCBI BioProject PRJNA1155696, BioSample SAMN43465610, with data under SRA accessions SRR31061614 and SRR31061614. Resequencing data for the M32 sample is under BioProject PRJNA1178157, BioSample SAMN44463567, and SRA accession SRR31129068.

## Acknowledgement

This work was partially supported by USDA-NIFA Hatch project COL00783A at Colorado State University.

