## Supplemental Figures and Tables for "*FHY3*/*FAR1* transposable elements generate adaptive genetic variation in the *Bassia scoparia* genome"

Supplemental table S1. Flow cytometry data estimating the genome size (1C) of *Bassia scoparia* to be ~900-1050 Mbp. A standard was run with each sample that was known to have 2.5 pg DNA/cell. Conversion of pg DNA to base pairs (978 Mbp/pg) derived from Dolezel et al (2003).

| Sample name | Sample ID | Standard(int.) G0+G1 mean | DNA content (pg/2C) | St. Dev ± |
|---|---|---|---|---|
| 7710 A | ***1*** | 257.39 | 2.17 | |
| | | 278.10 | 2.18 | |
| | | 301.78 | 2.16 | |
| | | 325.80 | 2.17 | |
| | | | **2.17** | **0.006** |
| 7710 B | ***2*** | 260.82 | 2.20 | |
| | | 282.69 | 2.18 | |
| | | 306.18 | 2.18 | |
| | | 329.44 | 2.18 | |
| | | | **2.19** | **0.008** |
| 7710 C | ***3*** | 252.69 | 2.17 | |
| | | 273.93 | 2.15 | |
| | | 295.06 | 2.16 | |
| | | 319.74 | 2.14 | |
| | | | **2.15** | **0.014** |
| 7710 D | ***4*** | 258.99 | 2.17 | |
| | | 281.12 | 2.16 | |
| | | 304.88 | 2.15 | |
| | | 327.03 | 2.17 | |
| | | | **2.16** | **0.007** |
| 7710 E | ***5*** | 252.11 | 2.17 | |
| | | 272.60 | 2.15 | |
| | | 294.68 | 2.15 | |
| | | 319.06 | 2.14 | |
| | | | **2.15** | **0.009** |
| 7710 F | ***6*** | 254.06 | 2.13 | |
| | | 273.54 | 2.13 | |
| | | 293.94 | 2.13 | |
| | | 316.73 | 2.14 | |
| | | | **2.13** | **0.008** |
| M32 A | ***10*** | 261.21 | 2.03 | |
| | | 280.72 | 2.03 | |
| | | 303.88 | 2.03 | |
| | | 328.30 | 2.03 | |
| | | | **2.03** | **0.004** |
| M32 B | ***11*** | 259.50 | 2.01 | |
| | | 281.57 | 1.98 | |
| | | 304.90 | 1.98 | |
| | | 328.47 | 1.98 | |
| | | | **1.99** | **0.013** |
| M32 C | ***12*** | 258.59 | 2.02 | |
| | | 278.88 | 2.01 | |
| | | 300.99 | 2.00 | |
| | | 327.10 | 2.00 | |
| | | | **2.01** | **0.007** |

Supplemental figure S1. Mitotic chromosome spread showing the 9 chromosome pairs of Bassia scoparia.

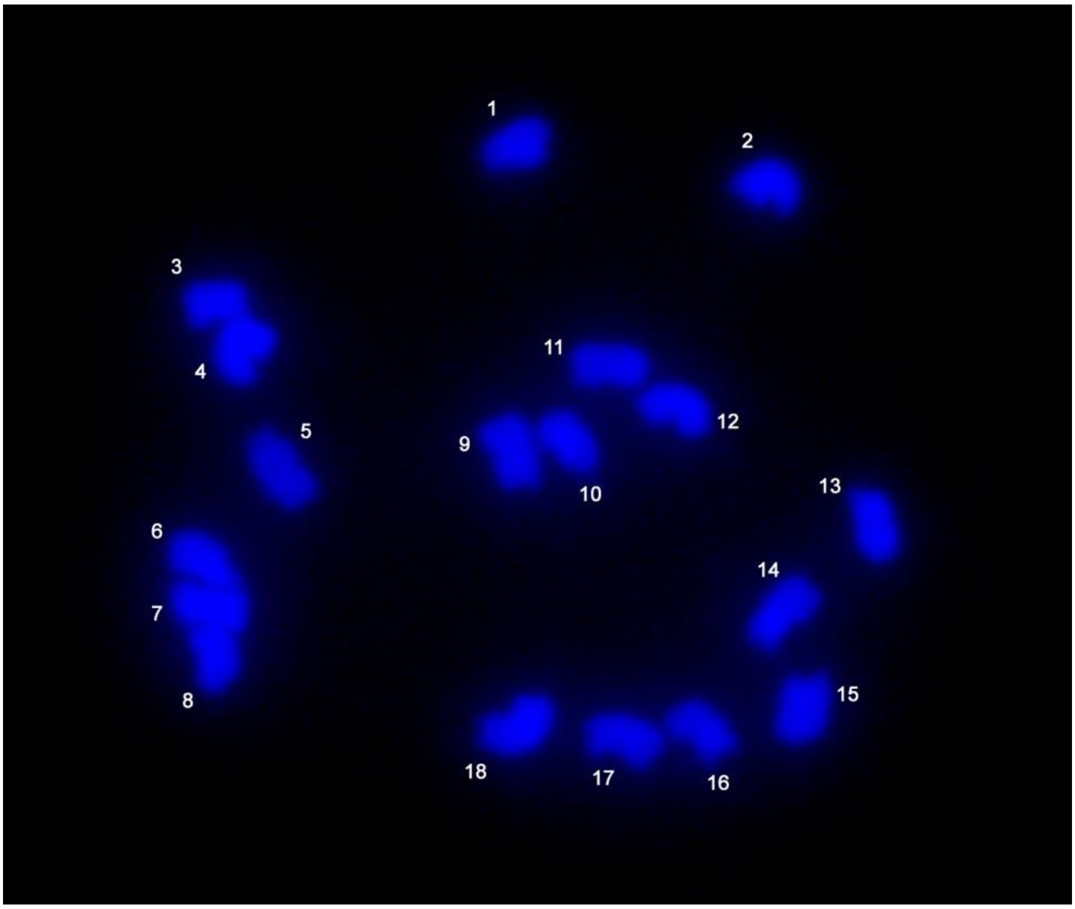

Supplemental Figure S2. Dotplot showing synteny between genome assemblies of *Bassia scoparia* (X-axis) and *Beta vulgaris* (Y-axis). Color represents number of orthologous genes in a 12-gene sliding window (blue=1, yellow =5+)

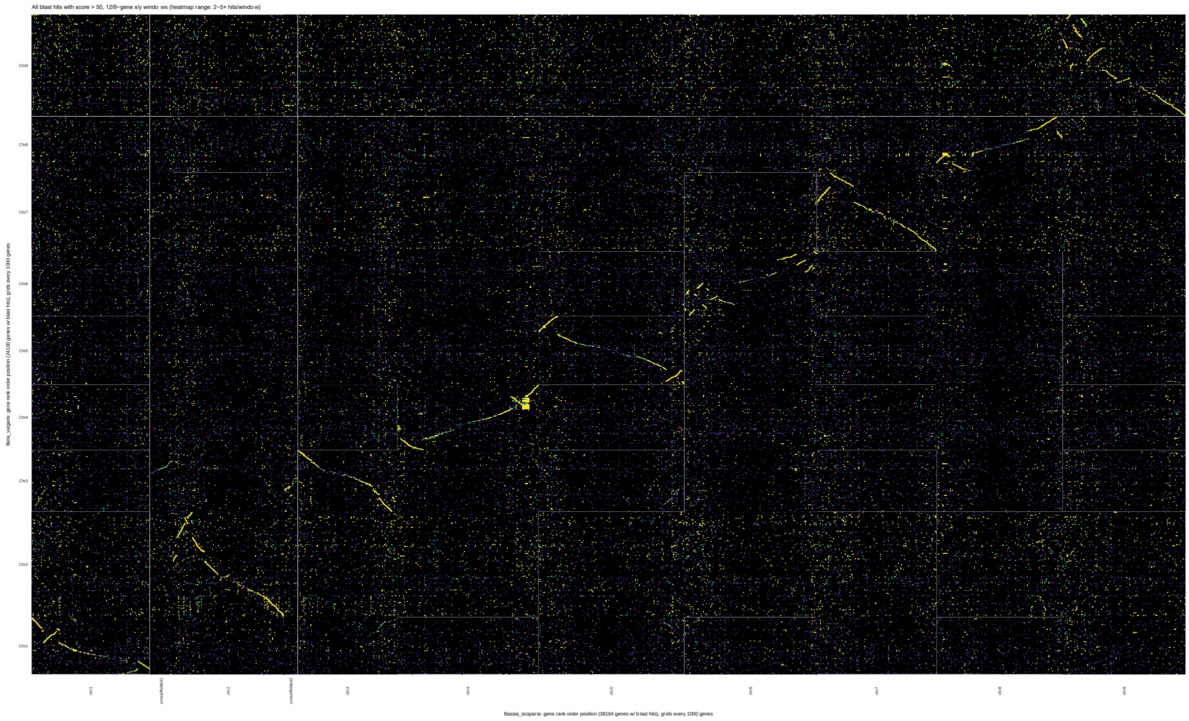

Supplemental SX: Number (and percent) of Benchmarking Universal Single Copy Orthologs (BUSCOs) from the embryophyte_obd10 dataset identified in either the *Bassia scoparia* genome assembly or annotated gene set.

| Dataset | Complete | Complete and single | Complete and duplicated | Fragmented | Missing |
|---|---|---|---|---|---|
| Genome | 1533 (95%) | 1483 (92%) | 48 (3%) | 30 (1.9%) | 51 (3.1%) |
| Transcriptome (CDS) | 1435 (88.9%) | 1283 (79.5%) | 152 (9.4%) | 76 (4.7%) | 103 (6.4%) |

Supplemental table S3. Information of kochia MGE validation primers.

| Assay | Primer name | Primer sequence (5' to 3') | Target length (bp) | Annealing temperature (°C) | Function |
|---|---|---|---|---|---|
| 1 | F1<br>R1 | CATCTTCTCTTCTCAACCACTC<br>ACCACACCCACTCATTCAC | 904 | 55 | Test for the presence of MGE |
| 2 | F1<br>R2 | CATCTTCTCTTCTCAACCACTC<br>CGGAAAACAACACTTAGAATGC | 403 | | Amplify upstream of MGE insertion site |
| 3 | F2<br>R3 | GTGAATGAGTGGGTGTGGT<br>TGAACAACTTAGCAGCAAGG | 222 | | Amplify downstream of MGE insertion site |
| 4 | F1<br>R4 | CATCTTCTCTTCTCAACCACTC<br>CTACAGTCAACAAAACCAAACC | 993 | | Amplify the upstream junction of MGE insertion |
| 5 | F3<br>R3 | ACCCAAATTCTTCAACAGCAC<br>TGAACAACTTAGCAGCAAGG | 931 | | Amplify the downstream junction of MGE insertion |

Supplemental Table S4. Summary of repetitive element content in the genome of *Bassia scoparia* as annotated by EDTA (Ou et al., 2019).

| Class | Count | Total Length (bp) | % of the genome |
|---|---|---|---|
| LINE | | | |
| Unknown | 2713 | 1515802 | 0.18 |
| LTR | | | |
| Copia | 84520 | 82902502 | 9.90 |
| Gypsy | 171846 | 196752767 | 23.49 |
| Unknown | 132386 | 87260301 | 10.42 |
| TIR | | | |
| CACTA | 76828 | 31975832 | 3.82 |
| Mutator | 111880 | 46921227 | 5.60 |
| PIF Harbinger | 28568 | 12055050 | 1.44 |
| Tc1 Mariner | 117029 | 31368013 | 3.75 |
| hAT | 45644 | 16314299 | 1.95 |
| nonTIR | | | |
| helitron | 156834 | 60708457 | 7.25 |
| Repeat region | 67491 | 24539723 | 2.93 |
| Total | 995739 | 592313973 | 70.72 |

Supplemental Table S5. Data sources for proteomes used in Orthofinder and CAFÉ analysis.

| Species | Source |
|---|---|
| Amaranthus hybridus | CoGe: ID57427 |
| Amaranthus tuberculatus | CoGe: ID 56882 |
| Amaranthus palmeri | CoGe: ID 55760 |
| Amaranthus hypochondriacus | CoGe: ID 34733 |
| Atriplex hortensis | CoGe: ID 56906 |
| Beta vulgaris | Phytozome: ID 782 |
| Spinacia oleracea | spinachbase.org/ftp/genome/Monoe-Viroflay/ |
| Chenopodium quinoa | CoGe: ID 60716 |
